# A Hybrid Learning Method with High Specificity for Excluding Non-chromosome Objects Problem

**DOI:** 10.1101/2022.10.06.511091

**Authors:** E-Fong Kao, Ya-Ju Hsieh, Chien-Chih Ke, Wan-Chi Lin, Fang-Yu Ou Yang, Jain-Shing Wu

**Author notes:** Address correspondence to: Jain-Shing Wu, Computer Science and Information Management, Providence University, Taichung City 433303, Taiwan, Phone: (Area Code 011-886-04) 2632-8001 ext. 18118, Fax: (Area Code 011-886-04) 2632-4045.

## Abstract

**Background:** For automated analysis of metaphase chromosome images, the chromosome objects on the images need to be segmented in advance. However, the segmentation results often contain a lot of non-chromosome objects in the images. Hence, elimination of non-chromosome objects is an essential process in automated chromosome image analysis. This study aims to exclude non-chromosome objects and preserve as many chromosomes as possible. In this paper, we propose a hybrid deep learning method to exclude non-chromosome objects from metaphase chromosome images.

**Method:** The proposed method consists of two phases. In the first phase, two classification results are obtained from feature-based and image-based convolutional neural networks (CNN) separately (the feature-based CNN uses the features of the images as input; the image-based CNN uses the images as input directly). In the second phase, the prediction results from the above two CNNs are combined and resent to another CNN to obtain final classification results.

**Results:** The proposed method uses 18,757 non-chromosome objects and 43,398 chromosomes (including single and multiple overlapped chromosomes) from 1038 chromosome images to evaluate the performance. The experimental results show that the proposed method can detect and exclude 99.61% (18,683/18,757) of the non-chromosome objects and preserve 99.95% (43,375/43,398) of the chromosomes for further analysis.

**Conclusions:** The proposed method has a high effectiveness on excluding non-chromosome objects and could be used as a preprocessing procedure for chromosome image analysis.

Graphic Abstract

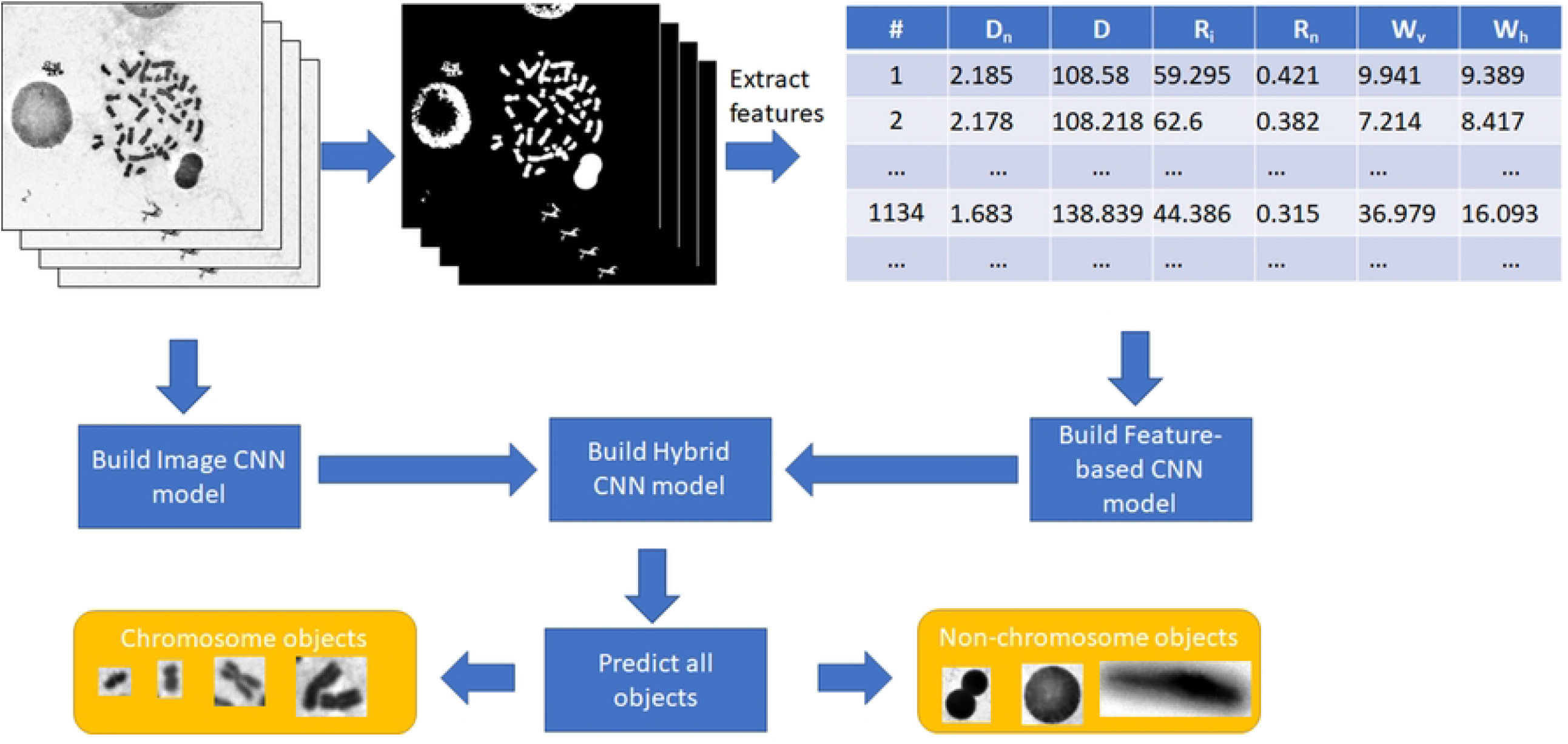

## 1. Introduction

For automated analysis of metaphase chromosome images, the chromosomes on the images need to be segmented first by performing thresholding [1–6] or other techniques [7–9]. Meanwhile, the segmentation results often contain several non-chromosome objects, such as interphase cells, stain debris, and other background noise. These non-chromosome objects need to be excluded from the segmented chromosome candidates for further analysis. Hence, excluding non-chromosome objects is an early step in automated chromosome image analysis.

The success of excluding non-chromosome objects has a large impact on the outcome of chromosome image analysis, such as chromosome counting [10, 11]. Eliminating non-chromosome objects is essential in chromosome image analysis, and studies on chromosome image analysis employed different methods to achieve this goal. Ji et al. [2] proposed a fully automatic chromosome segmentation method involving a rule-based approach to eliminate non-chromosome objects. Wang et al. [11] proposed a fully-automated computer-aided method to detect metaphase chromosomes, count the total number of chromosomes, and compute the DNA index. In the method, a knowledge-based classifier based on the seven features is applied to delete non-chromosome objects. Uttamatanin et al. [12] proposed a metaphase selection tool that classifies segmented objects into four classes based on width, height, and estimated area ratio. One (Class 4) of the four classified classes corresponds to non-chromosome objects. Sayed et al. [13] proposed a fully automatic approach based on Fast Fuzzy C-Means and Grey Wolf Optimization to remove interphase cells and extract chromosomes from metaphase chromosome images. Li et al. [14] proposed a method for the automated discrimination of dicentric and monocentric chromosomes. In the method, the non-chromosome objects are filtered by examining the sizes, brightness, and contours after the segmentation of all objects in an image.

Although related studies employed different methods to eliminate non-chromosome objects, preserving as many chromosomes as possible to exclude non-chromosome objects is still a worthwhile issue.

Nowadays, the deep learning methods, such as Convolution neural network (CNN) [15], are adopted for classification problem and obtained very good performance. Priya et al. [16] use Residual network (ResNet) to predict the COVID-19. They use different layer compositions to handle with lung CT images. Faruqui et al. [17] using CNN to classify the lung cancer from CT and wearable sensor-based medical IoT (MIoT) data. Masood et al. use deep fully convolutional neural network to lung cancer. They also compared their results with CNN model. Their model gets higher accuracies with 6 different datasets [18]. Zhou et al. use CNNs via the status lymph node metastasis to predict the primary breast cancer [19]. In their results, they obtained high AUC results for final clinical diagnosis. Kuntz et al. [20] review the deep learning methods adopted for gastrointestinal cancers. They point out that the CNN-based methods are good to predict gastrointestinal cancers but requiring for large number of clinical trials to support.

Based on the efficiency and powerfulness of the deep learning methods, we propose a hybrid deep learning method to exclude non-chromosome objects in metaphase chromosome images. The method consists of two phases. In the first phase, two classification results are obtained from different CNNs separately (one uses the features of the images as input; another uses the images as input directly). In the second phase, the classification results obtained in the first phase are combined and resent to another CNN for further classification. At final, the effectiveness of the proposed method is evaluated in this study.

## 2. Materials and methods

In this paper, we propose a hybrid method based on the deep learning in two phases. The flowchart of the proposed method is shown in Fig. 1(a). In the first phase, the input data (images) are inputted to the two separated ways, Image CNN way and feature extraction way. In Image CNN way, the cropped images are regarded as input and directly used for CNN classification. Details of the Image CNN is shown as Fig. 1(b). In the feature extraction way shown in Fig. 1(c), the features of the segmented objects in the images are extracted and used for classification with another Convolutional Netural Network. In the second phase, all prediction results of all objects from Image CNN and Feature-based CNN are combined together for further classification. And then, the combined results are sent to the Hybrid CNN to obtain final classification results. All the details are described as follows:

**Fig. 1.**
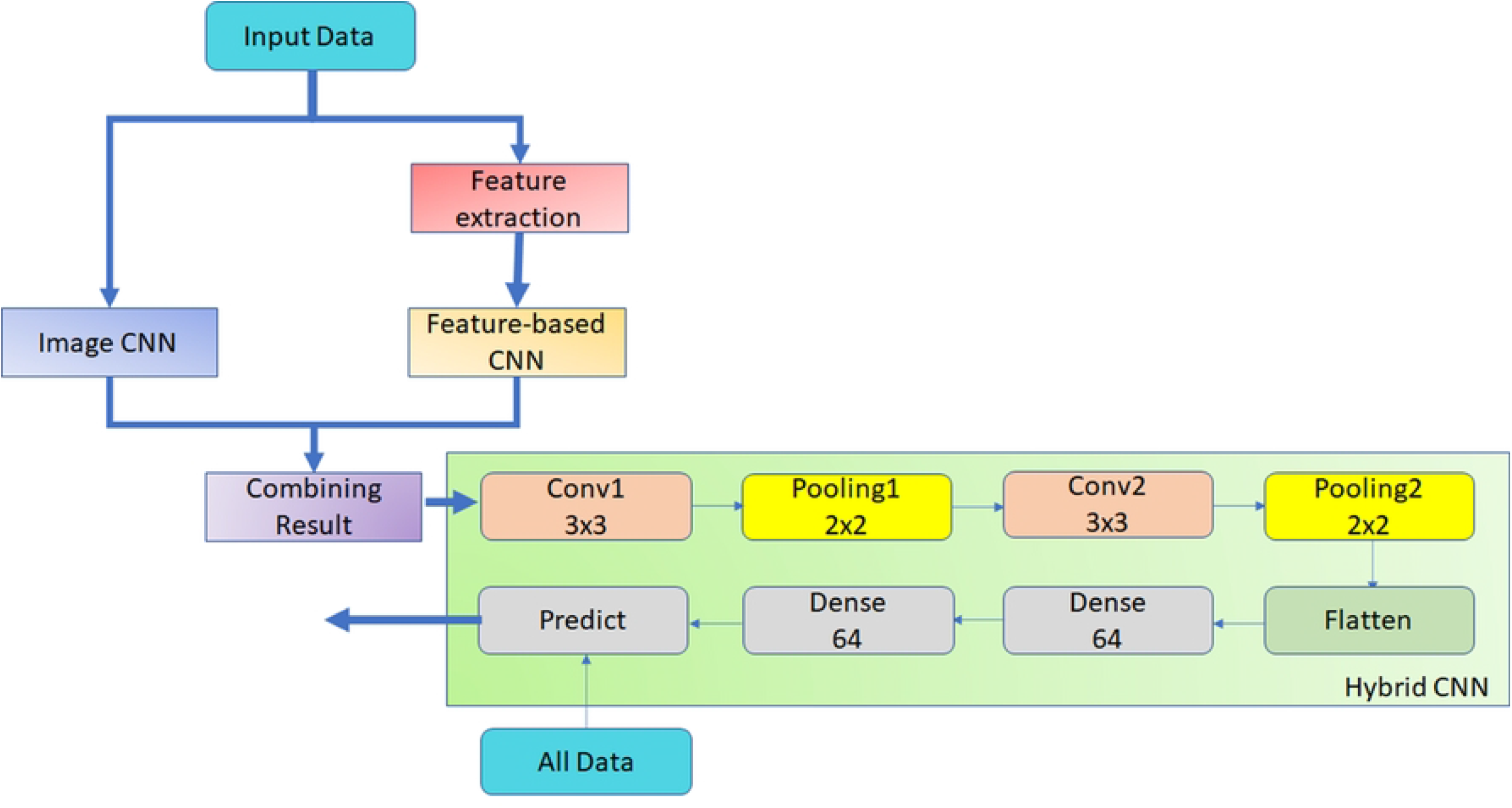
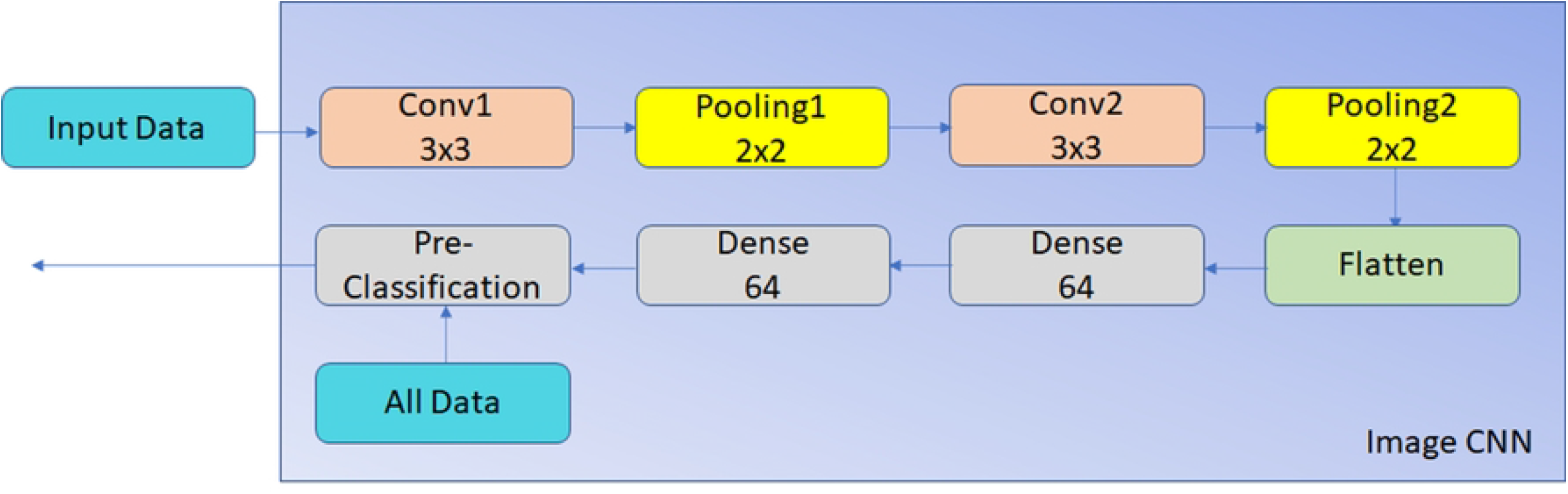
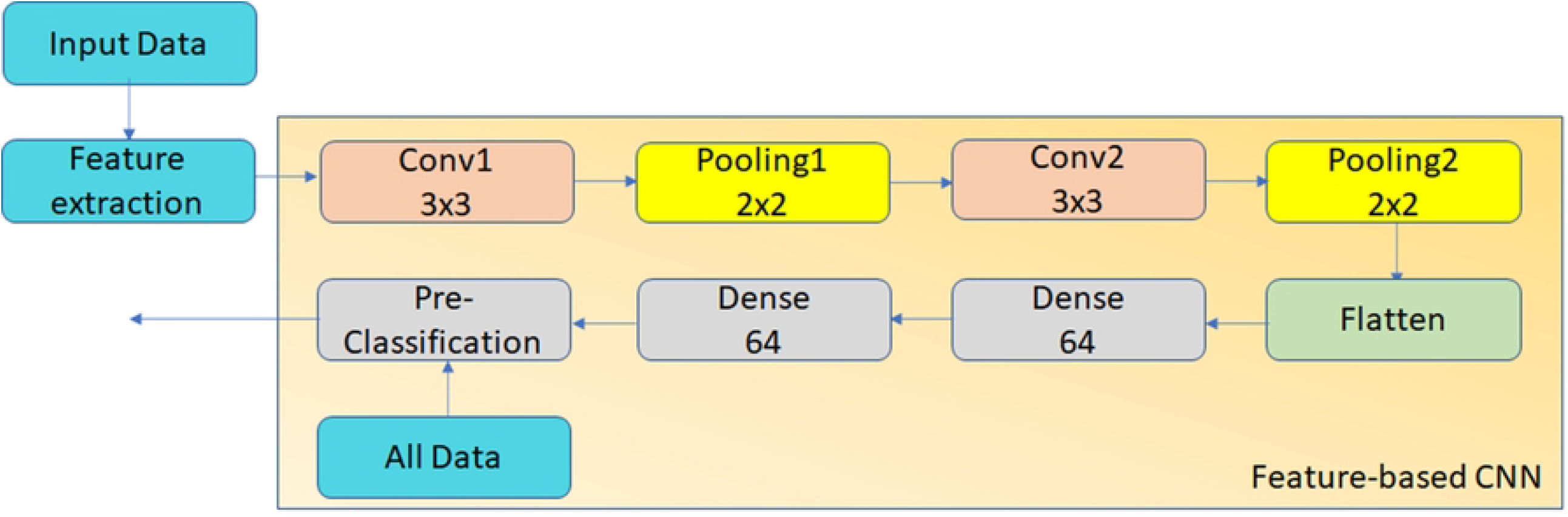
The flowchart of the proposed method. (a) the whole flowchart of the Hybrid CNN. (b) the flowchart of Image CNN. (c) the flowchart of Feature-based CNN.

### 2.1. Image Database (Data source)

The proposed method is evaluated using 1038 metaphase chromosome images obtained from the Institute of Nuclear Energy Research in Taiwan. All chromosome images are color JPEG format obtained using a microscope system (Zeiss, Imager Z2) with an objective lens of 64× magnification and a scanning system (Metasystems, Metafer4) with the image size of 1360×1024 pixels. A color JPEG image comprises red, green and blue components. Each pixel value (R, G, B) in color JPEG images is converted to a gray scale value between 0 and 255 for further analysis. The higher and lower pixel values correspond to the background (bright regions) and foreground (dark regions) in the images, respectively.

### 2.2. Chromosome candidate segmentation

A chromosome image and its histogram in terms of gray scale value are shown in Figs. 2(a) and 2(b), respectively. The histogram has two major clusters; one with the lower value corresponds to the chromosomes or the foreground, and the other with the higher value corresponds to the background. A threshold between the two clusters must be determined in advance to segment the chromosomes. In this study, the threshold is determined by the mean value of two clusters, as shown in Fig. 2(b). For the chromosomes, a gray scale value with maximum number *Peak*_*fore*_ is searched between 0 and 100; then, *Peak*_*bkg*_ is searched between 150 and 255 for the background. The threshold is calculated as (*Peak*_*fore*_+*Peak*_*bkg*_)/2. A binary image for the chromosome candidates is obtained by segmenting the pixels with values less than the threshold, as shown in Fig. 2(c). These segmented objects include chromosomes and non-chromosome objects. Excluding non-chromosome objects and preserving as many chromosomes as possible are the main goals of this study.

**Fig. 2.**
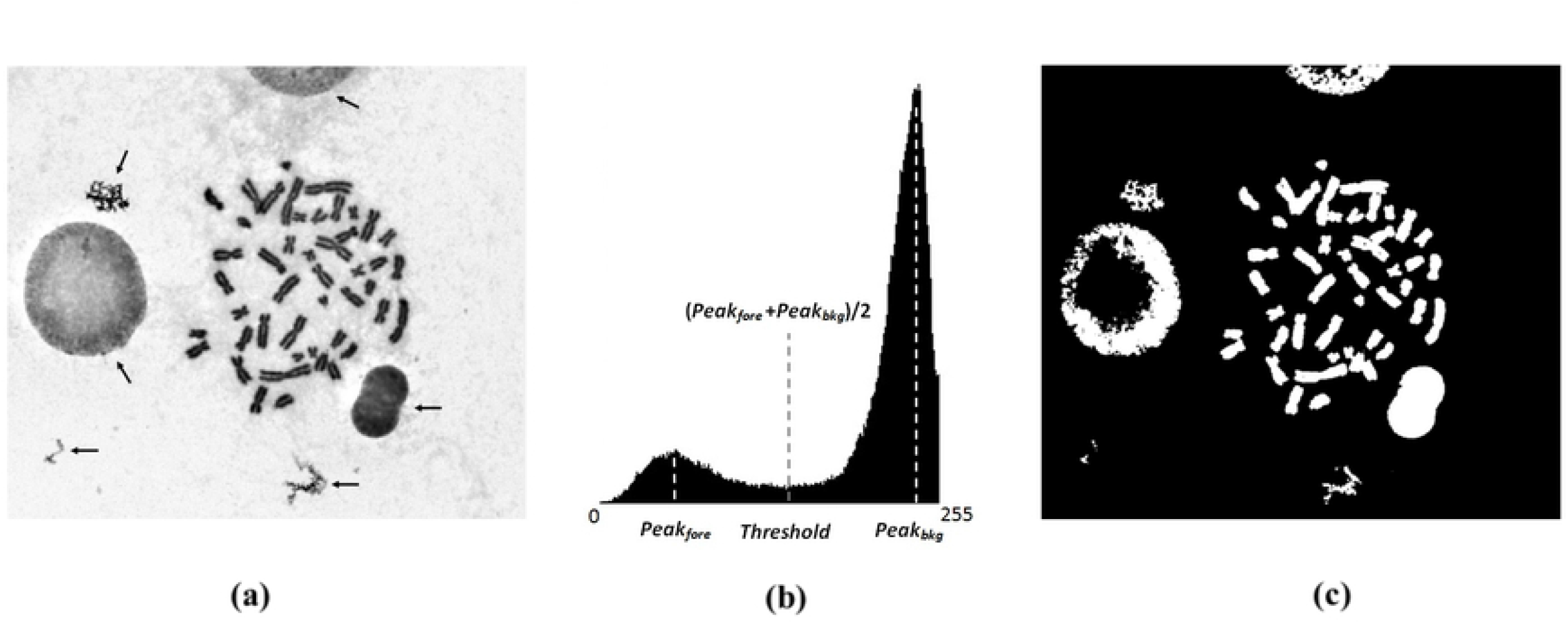
(a) Chromosome image with non-chromosome objects indicated by the arrows; (b) histogram of Fig. 2(a) and threshold used to segment the chromosome candidates; and (c) segmented objects corresponding to Fig. 2(a).

### 2.3. Image Convolutional Neural Network (Image CNN)

As shown in the Fig. 1(b), the data (images) are directly used for CNN classification. Subimages containing the corresponding segmented objects described in section 2.2 are cropped from original metaphase chromosome images. All the cropped images are used as input, and sent to the CNN. Here we use the 3×3 Convolution filters in convolution layer and 2×2 max pooing filters in Pooling layer. After twice convolution and pooling layers, the results are flatten and sent to full connected neural networks which consisted two layers and each layer has 64 units.

Note that the numbers of the cropped images are large, hence, we only randomly choose 40000 images as training dataset to train the model. After training process, the whole images are sent to the model to predict the results. The final probabilities of all objects are kept for further classification.

### 2.4. Feature-based Convolutional Neural Network (Feature-based CNN)

After segmenting the chromosome candidates, four classes of features, namely, area, density-based features, roughness-based features, and widths, of the segmented candidates are extracted for CNN to discriminate between chromosomes and non-chromosome objects. After the features are extracted, these feature values are sent to CNN to learn the model. The methods to measure the features are described in detail as follows.

#### 2.4.1. Area

The area is an important feature for the differentiation between chromosomes and non-chromosome objects, as shown in Fig. 2(a). The areas of chromosomes fall in a specific range; otherwise, the areas of non-chromosome objects vary widely. The area (total number of the pixels) of each segmented candidate is used as one feature for the differentiation between chromosomes and non-chromosome objects.

#### 2.4.2. Density-based features

As shown in Fig. 2(a), the chromosomes represent highly dense objects in the image. Density is another important feature of chromosomes. Hence, the average density value of each segmented candidate is used as a feature to differentiate chromosomes from non-chromosome objects. Assuming that the number of pixels of a segmented object is *n*, the density function for the segmented object can be defined as *Gray(i)*, where *i* = *1, 2*, …, *n*. The average density, *D*, of the segmented object is formulated as follows:

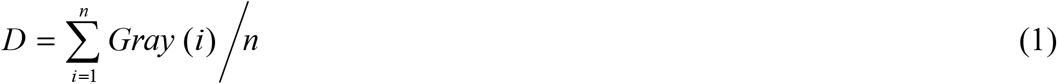

The density values may vary between the chromosome images because of different imaging conditions, as shown in Figs. 3(a) and 3(b). A normalized density-based feature is used to reduce the impact of different imaging conditions. For this purpose, the mean density of the chromosomes in each chromosome image is obtained for normalization. Assuming that most chromosomes would locate centrally in the image, the segmented objects in the rectangular region located centrally are used to determine the mean chromosome density for each image, as shown in Fig. 3(c). The mean chromosome density for each image *Mean*_*chromosome*_ can be obtained simply by the sum of the density values of all segmented pixels divided by the total number of segmented pixels in the rectangle. To use *Mean*_*chromosome*_ as the normalized factor for reducing the influence of the density variation between the images, the density *D* of each segmented object is normalized to *Mean*_*chromosome*_. The normalized density *D*_*n*_ can be obtained for each segmented object by using Eq. (2):

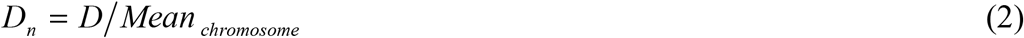

**Fig. 3.**
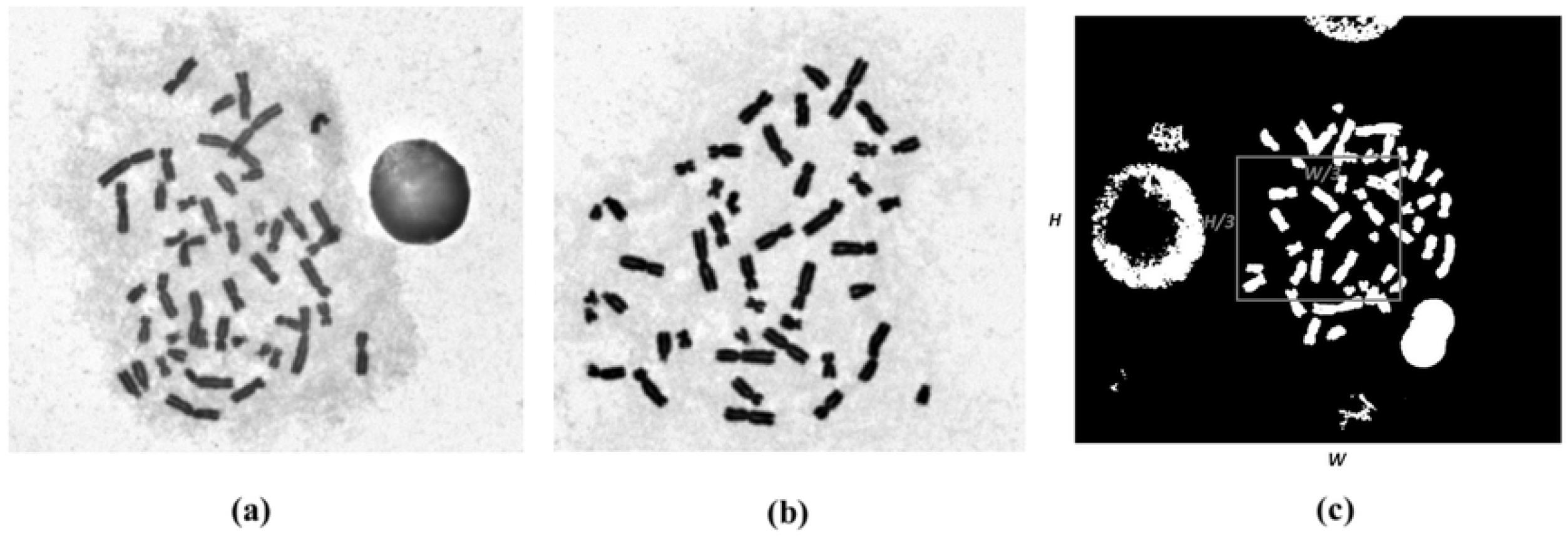
(a) Low-contrast and (b) high-contrast chromosome images due to different imaging conditions (c) rectangular region in which the segmented objects were used to determine the normalized factor.

Density-based features *D* and *D*_*n*_ are used to exclude non-chromosome objects with either higher or lower density than chromosomes.

#### 2.4.3. Roughness-based features

Roughness is an obvious feature that differs between chromosomes and some non-chromosome objects, as shown in Figs. 4(a) and 4(c). Fig. 4(a) shows that the roughness of a non-chromosome object is considerably higher than that of a chromosome. By contrast, as shown in Fig. 4(c), non-chromosome objects represent homogeneous or clearness inside, and the roughness inside the objects would be relatively lower than those inside chromosomes.

**Fig. 4.**
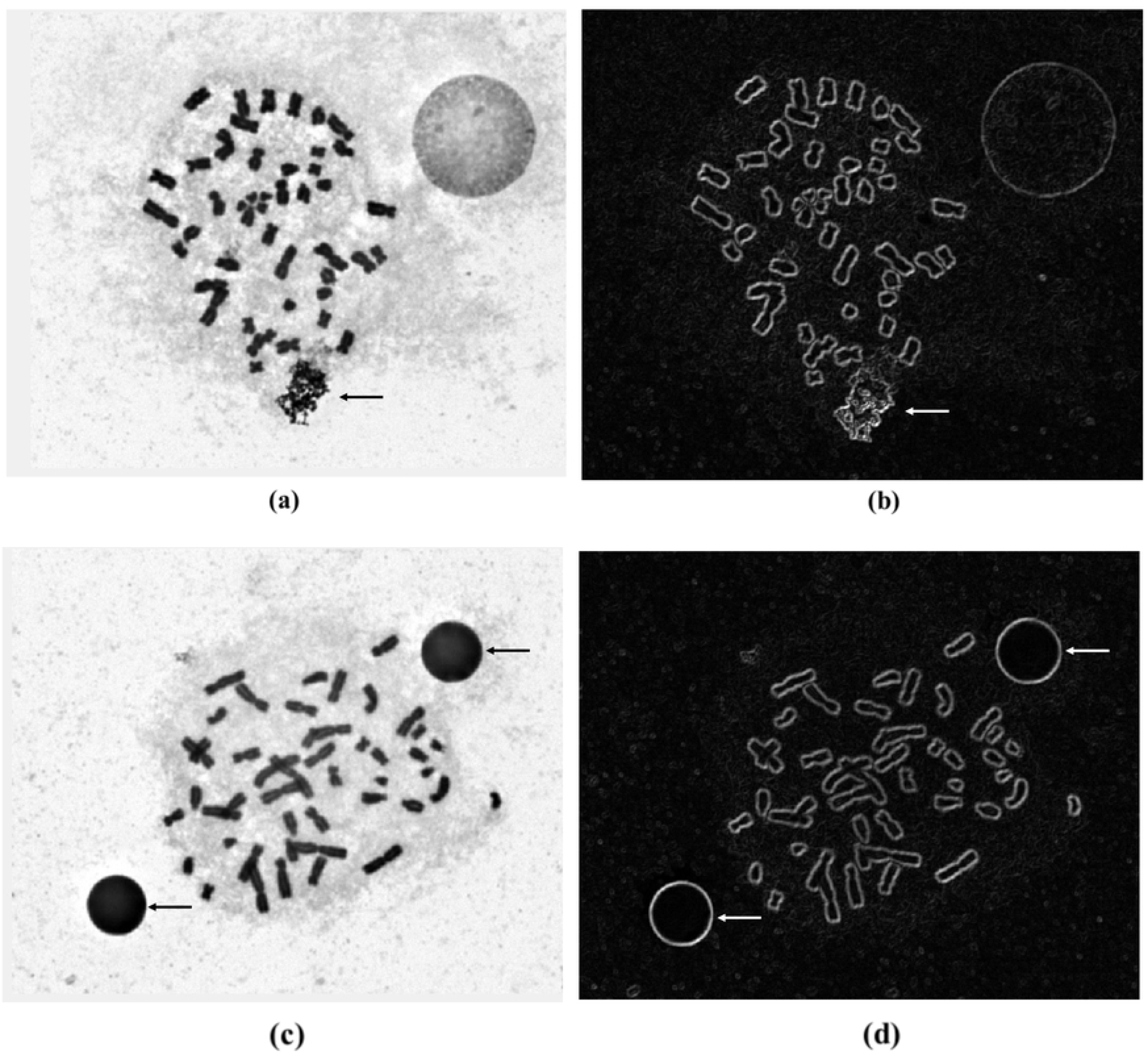
(a) Chromosome image with high-roughness non-chromosome object indicated by the arrow and (b) its corresponding gradient image. (c) Chromosome image with low-roughness non-chromosome objects and (d) its corresponding gradient image.

In this study, the gradient is used to measure the roughness of segmented objects.

The gradient value for each pixel in a segmented object is obtained by gradient operation [21], as shown in Fig. 5. For each pixel Z_5_ of which the gradient value is determined, the derivatives in the *x* and *y* directions are first obtained by applying the masks in Figs. 5(b) and 5(c), respectively. The corresponding equations are as follows:

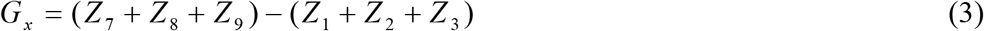

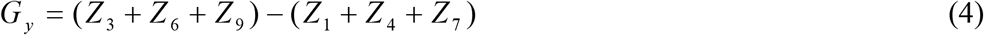

where Z_1_, Z_2_, Z_3_, Z_4_, Z_6_, Z_7_, Z_8_, and Z_9_, are the gray scale values of neighbor pixels around point Z_5_. Subsequently, the gradient value is determined by summing the absolute derivatives in the *x* and *y* directions as follows:

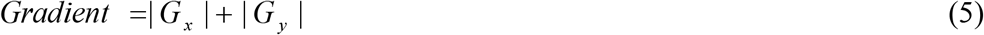

**Fig. 5.**
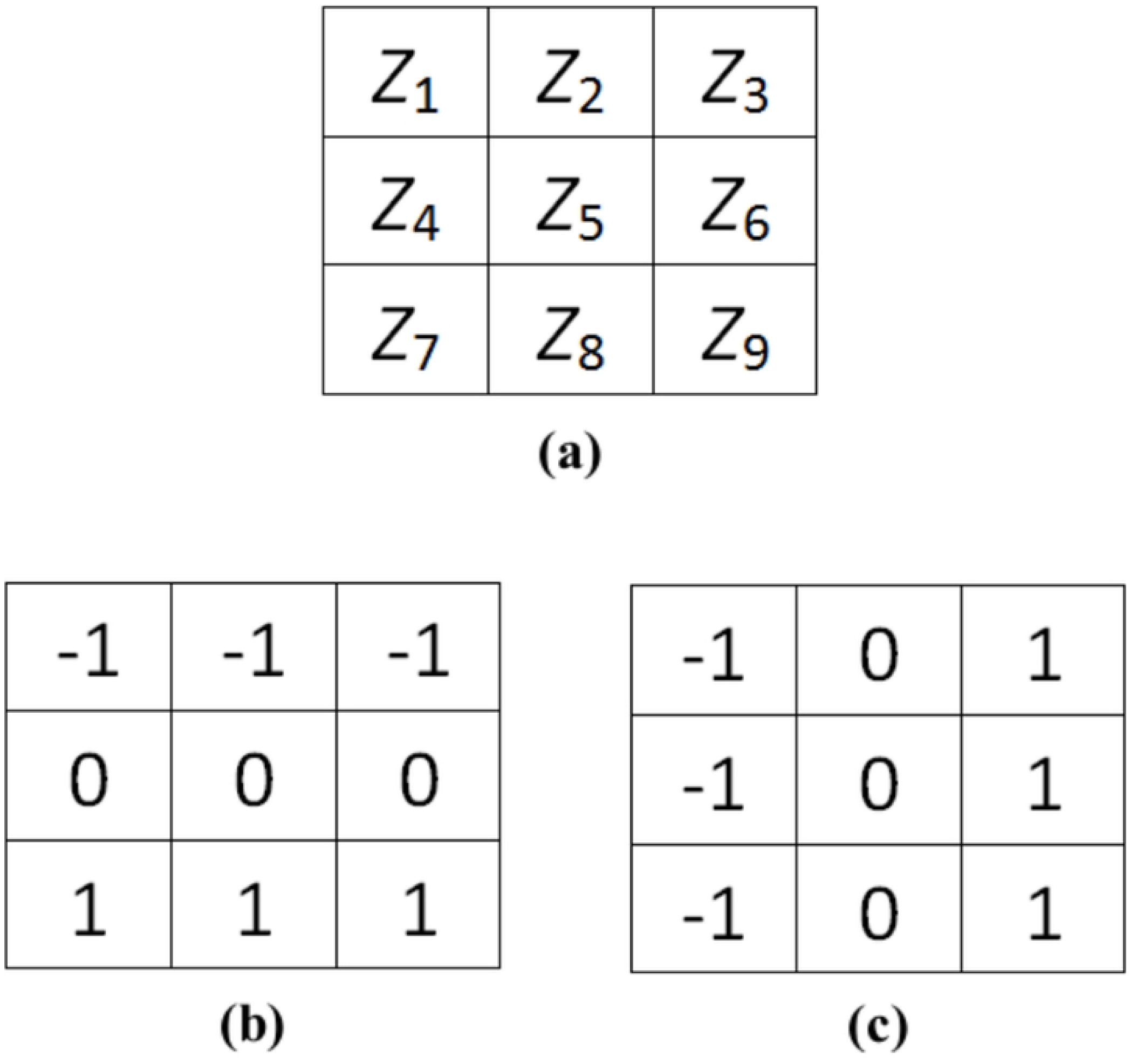
(a) A 3 × 3 region of an image and the masks used to compute the derivatives in (b) the *x* direction and (c) the *y* direction at point *Z*_5_.

Figures 4(b) and 4(d) show the gradient images corresponding to Figs. 4(a) and 4(c), respectively. For the non-chromosome object with high roughness, as shown in Fig. 4(b), the whole object represents high gradient value. By contrast, only the boundary represents high gradient value for chromosomes. For non-chromosome objects with low roughness inside, as shown in Fig. 4(d), the inside of the object represents very low gradient value. The roughness indices for the whole and the inside of segmented objects are used to exclude non-chromosome objects with either high or low roughness.

Assuming that the number of pixels of a segmented object is *n*, the gradient function for the segmented object can be defined as *Gradient*(*i*), where *i* = *1, 2*, …, *n*. The average gradient of the segmented object is formulated as follows:

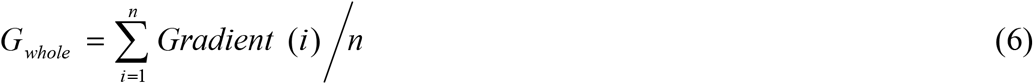

Similar to that described in Section 2.3.2, high- or low-contrast images caused by different imaging conditions also result in the variation of gradient value. The mean gradient value is used as the normalized factor for each chromosome image to overcome this problem. As shown in Fig. 3(c), the mean gradient value for each image *G*_*mean*_ can be obtained simply by the sum of the gradient values of all segmented pixels divided by the total number of segmented pixels in the rectangle. To use *G*_*mean*_ as the normalized factor for reducing the impact due to image contrast, the normalized roughness index, *R*_*n*_ can be obtained as follows:

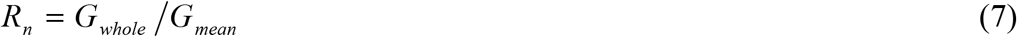

*R*_*n*_ is used to indicate the roughness of the whole segmented object.

An equivalent circle is obtained in advance to measure the roughness inside the segmented objects. Assuming the area of a segmented object is *A*, the radius *r* of the equivalent circle can be obtained by 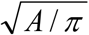. With the mass center of the segmented object as the center of the equivalent circle, the area covered by the equivalent circle is shown in Fig. 6. The inner circle with radius *r*/2 is used to measure the roughness inside the segmented object. Assuming that the number of pixels in the inner circle is *m*, the gradient function for the inner circle can be defined as *Gradient*_*inner*_(*j*), where *j* = *1, 2*, …, *m*. The roughness index *R*_*i*_ for the inner circle is formulated as follows:

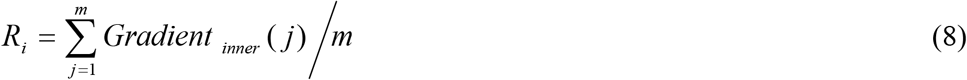

where *R*_*i*_ is the roughness inside the segmented object.

**Fig. 6.**
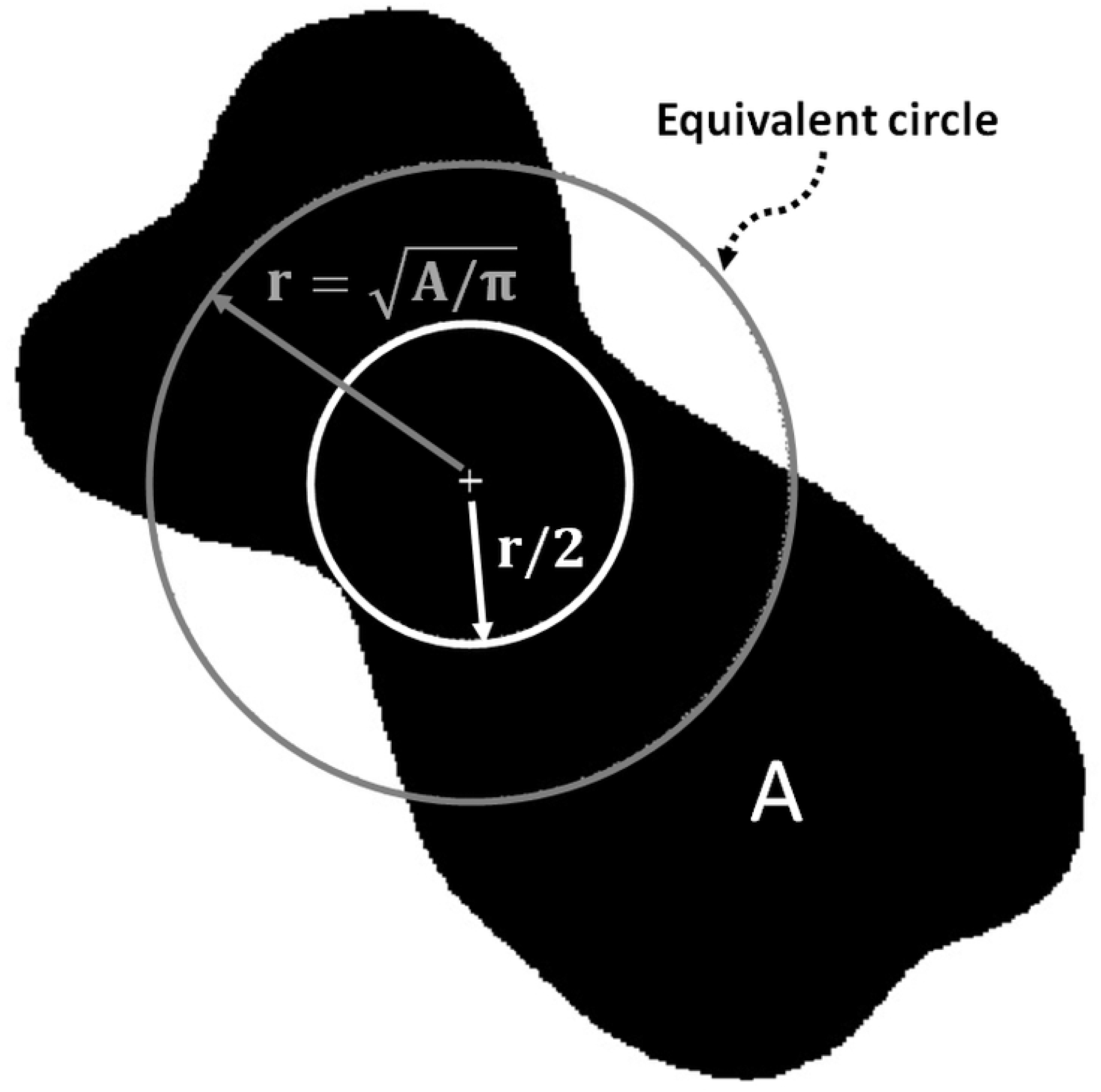
Equivalent circle and its inner circle used to compute the roughness index inside the segmented object.

The two roughness-based features *R*_*n*_ and *R*_*i*_ are used to exclude non-chromosome objects with either higher or lower roughness than chromosomes.

#### 2.4.4. Horizontal and vertical widths

Aside from the areas of chromosomes, their widths also fall in a specific range. Hence, non-chromosome objects with excessively large or small widths can be excluded on the basis of this range. Two features regarding segmented object width are used in this study. Figs. 7(a) and 7(b) represent the vertical and horizontal widths of the segmented object, respectively. The functions for vertical and horizontal widths are defined as *L*_*vertical*_(*i*) and *L*_*horizontal*_(*j*), respectively. The mean vertical width *W*_*v*_ and the mean horizontal width *W*_*h*_ of the segmented object are formulated as follows:

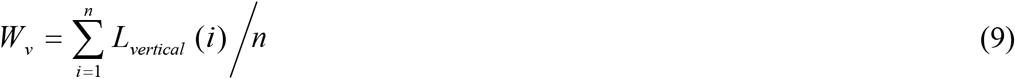

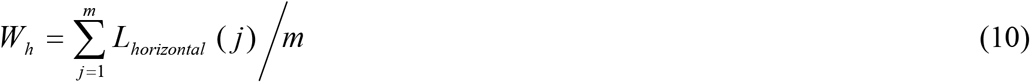

**Fig. 7.**
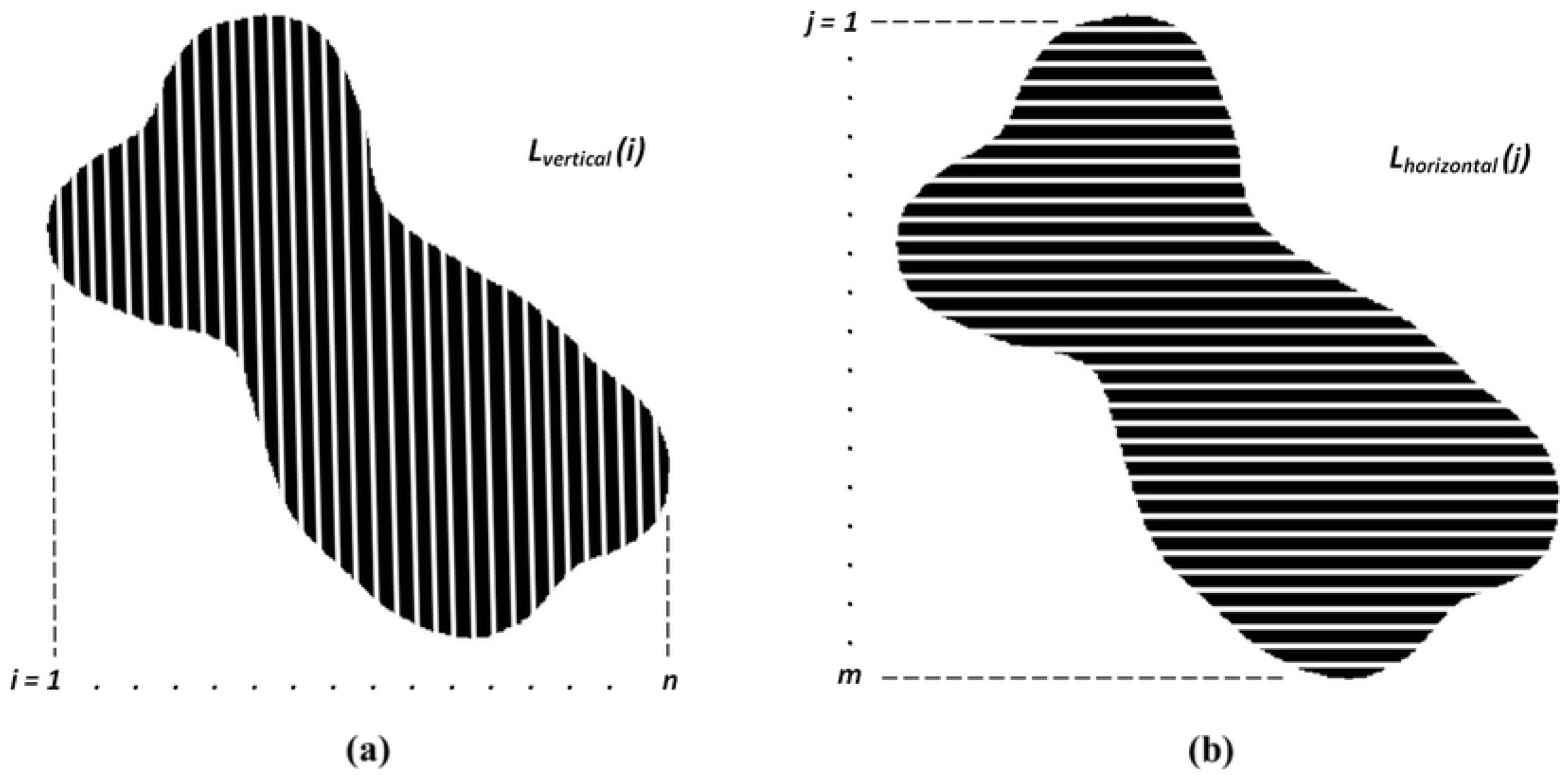
(a) Vertical width and (b) horizontal width of the segmented object.

The two width features *W*_*v*_ and *W*_*h*_ are combined to distinguish between chromosomes and non-chromosome objects.

#### 2.4.5. Classification with Convolutional Neural Network

After the features are extracted from the segmented objects, these features are sent to CNN to learn the model for distinguishing between chromosomes and non-chromosome objects. Similar to Section 2.3, the CNN we used is the same structures as shown in Fig. 1(c). The data first are inputted to the convolution layer. Here, we use “ReLU” as the activate function in all convolution layers. And then, the Max pooling is used in the Pooling layer. After twice convolution and pooling, the data are sent to flatten layer and for further full connected neural network to learn. After training and obtaining the model, the testing data are used to test the trained model. After building the model, the features of all segmented objects are sent to the model to predict the results. The measured probabilities of all segmented objects are obtained and sent to the Hybrid CNN for further classification.

### 2.5. Hybrid Convolutional Neural Network

In Fig. 1(a), after obtaining two CNN results, the prediction probabilities of each segmented object from Image CNN and Feature-based CNN are combined together for further classification. The combined results used as input are sent to Hybrid CNN to achieve higher classification performance. The Hybrid CNN structure is similar to Image CNN and Feature-based CNN. Also, 40000 data of segmented objects are randomly chosen as training dataset to train the model. After the model is built, the data of all the segmented objects are sent to the model to obtain final classification performance between chromosomes and non-chromosome objects.

## 3. Results

### 3.1. Usefulness of features

In Feature-based CNN, seven features are used to discriminate between chromosomes and non-chromosome objects. These features are evaluated in this section. Fig. 8 shows the distributions of chromosomes and non-chromosome objects in terms of the area in pixels. The results indicate that the areas of the chromosomes fall in a small specific range, but those of non-chromosome objects vary widely. The area of a segmented object is a significant feature for distinguishing between chromosomes and non-chromosome objects. Excessively small or large segmented objects should be considered as non-chromosome objects. Areas larger than 4000 pixels or smaller than 100 pixels are considered to indicate non-chromosome objects, and the remaining non-chromosome objects of which the areas are similar to chromosomes can be distinguished by other six features. Figs. 9, 10 and 11 show the distributions of all the chromosomes and remaining non-chromosome objects in terms of density-based features (*D* and *D*_*n*_), roughness-based features (*R*_*n*_ and *R*_*i*_) and widths (*W*_*h*_ and *W*_*v*_), respectively. The results indicate that these features have great effects on distinguishing between chromosomes and non-chromosome objects.

**Fig. 8.**
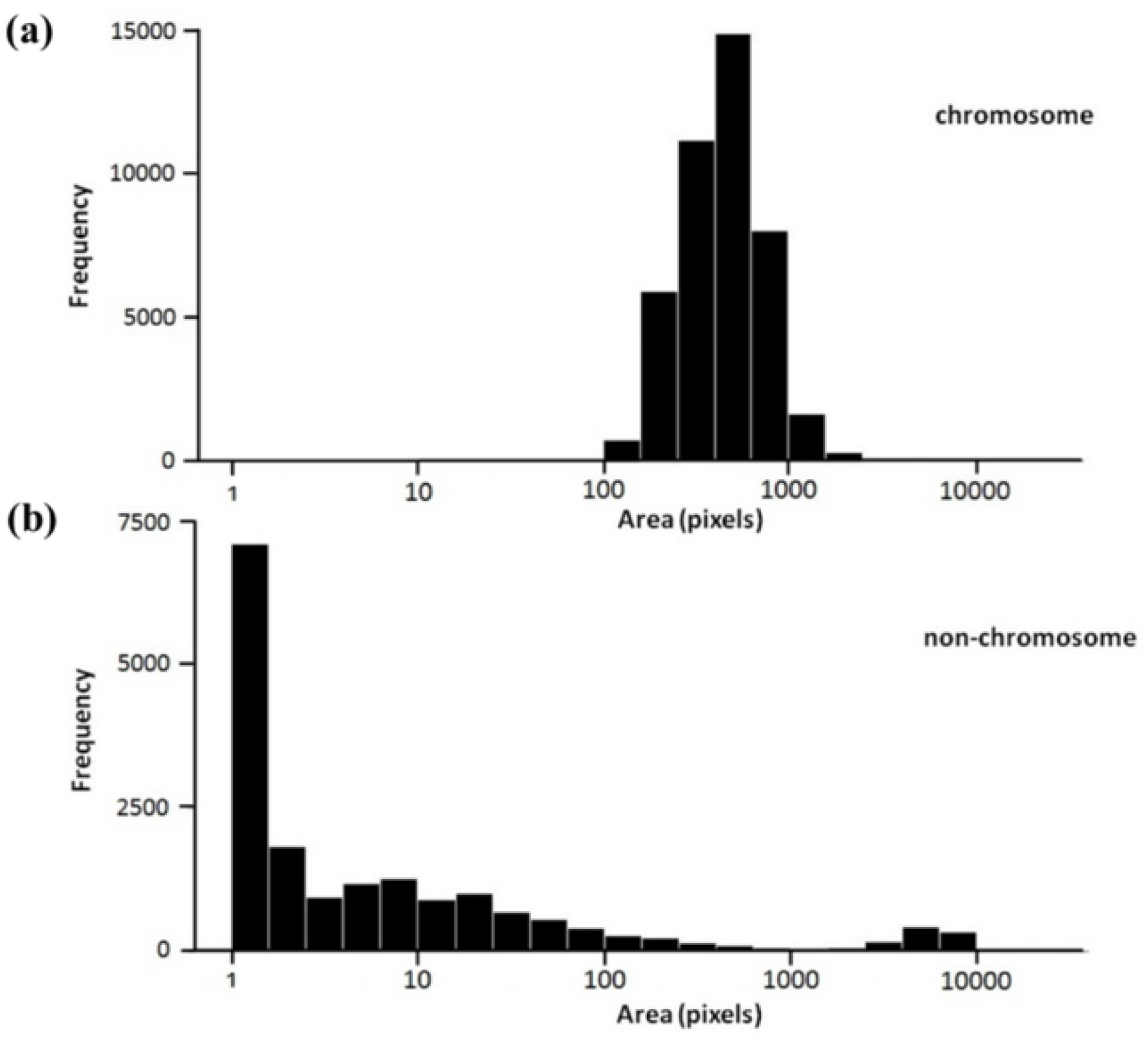
Distributions of (a) chromosomes and (b) non-chromosome objects in terms of area in logarithmic scale.

**Fig. 9.**
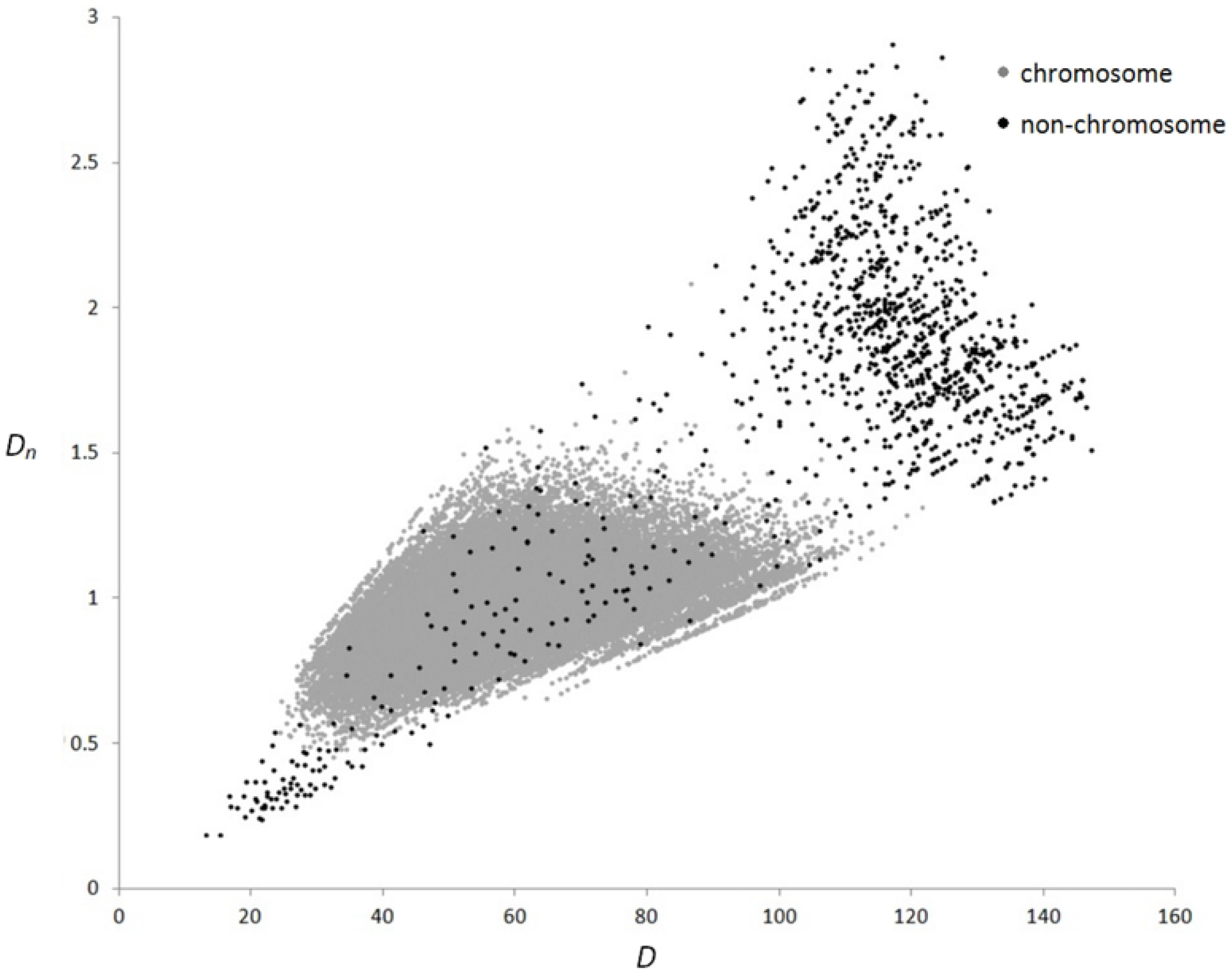
Distributions of all the chromosomes and remaining non-chromosome objects in terms of density-based features *D* and *D*_*n*_.

**Fig. 10.**
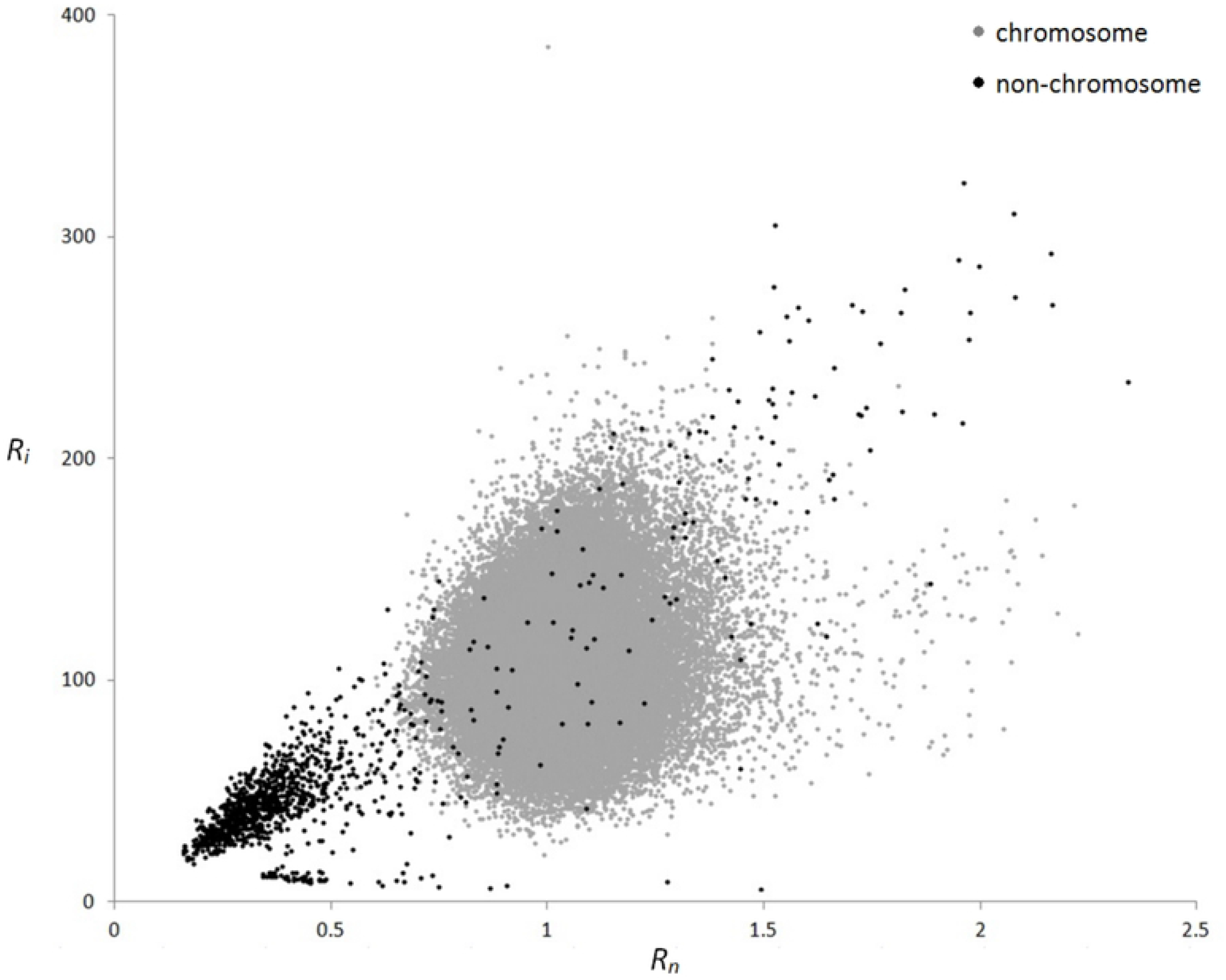
Distributions of all the chromosomes and remaining non-chromosome objects in terms of roughness-based features *R*_*n*_ and *R*_*i*_.

**Fig. 11.**
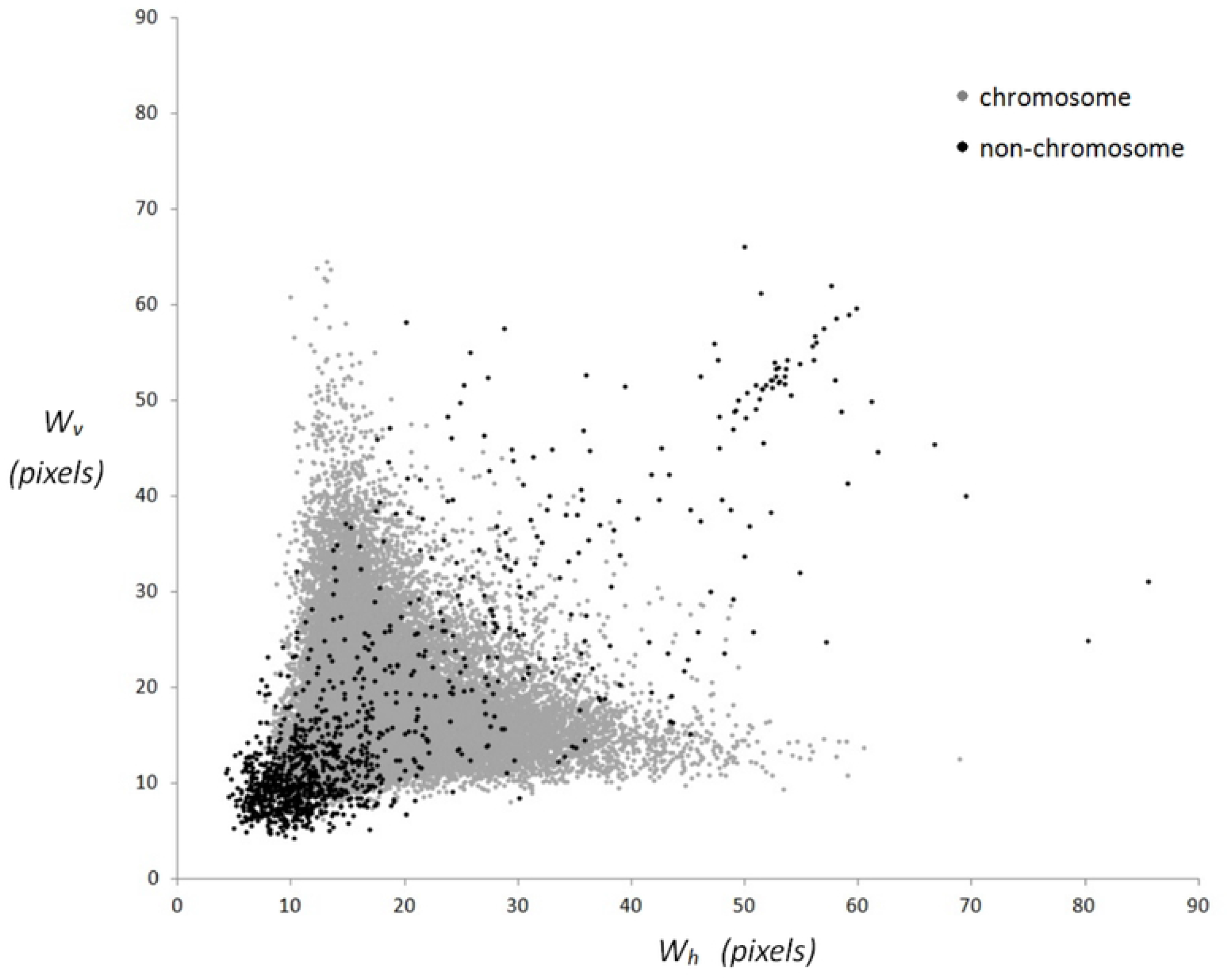
Distributions of all the chromosomes and remaining non-chromosome objects in terms of mean horizontal and vertical widths *W*_*h*_ and *W*_*v*_.

### 3.2. Classification results

All segmented objects in 1038 chromosome images, as described in Section 2.2, were classified as chromosomes or non-chromosome objects by two experienced experts. A total of 62,155 segmented objects were divided into 43,398 chromosomes (including single and multiple overlapped chromosomes) and 18,757 non-chromosome objects. This classified results are used as the gold standard to evaluate the proposed method.

Table 1 shows the classification results of Image CNN, Feature-based CNN and Hybrid CNN, respectively. Among three CNNs, Hybrid CNN has highest classification accuracy. The results indicate that the combination of Image CNN and Feature-based CNN can achieve higher classification performance further.

**Table 1.**
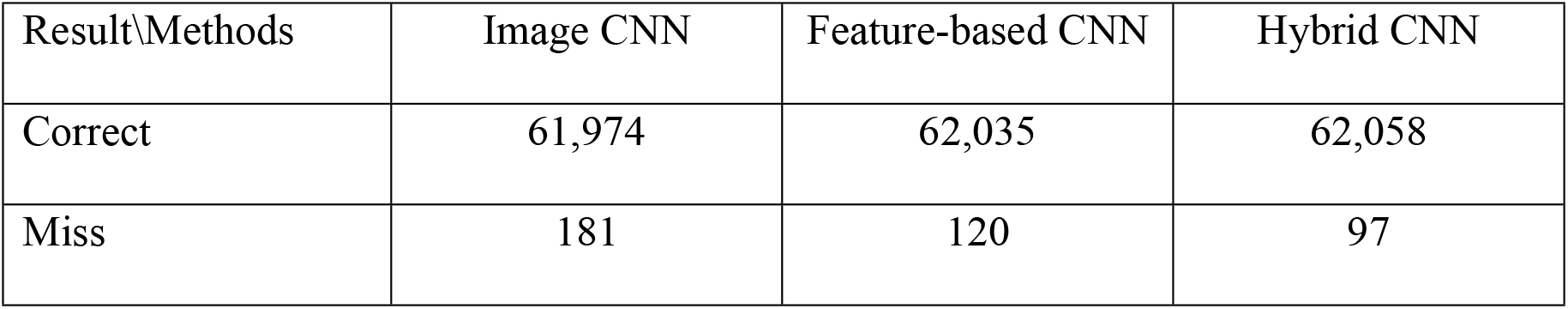
The comparison of classification results among Image CNN, Feature-based CNN and Hybrid CNN.

| Result/Methods | Image CNN | Feature-based CNN | Hybrid CNN |
| --- | --- | --- | --- |
| Correct | 61,974 | 62,035 | 62,058 |
| Miss | 181 | 120 | 97 |

For Hybrid CNN classification among 62,155 segmented objects, 62,058 objects are classified correctly, and only 97 objects are mis-classified. The classification accuracy is 99.84%. Among 97 mis-classified objects, there are 23 chromosome objects and 74 non-chromosome objects. The experimental results show that the proposed method can detect and exclude 99.61% (18,683/18,757) of the non-chromosome objects and preserve 99.95% (43,375/43,398) of the chromosomes for further analysis.

The experimental results show that the proposed method can really be used in excluding non-chromosome objects and determining the chromosome objects.

## 4. Discussion

In recent years, two main approaches are used to perform image recognition, image feature analysis method and CNN using images as input directly. In an image feature analysis method, the image features need to be designed manually and extracted from the images, and image recognition is performed by classification based on the extracted features. In addition, a CNN using images as input extracts or learns image features from an image via convolution and pooling process directly. In this study, we use both image feature analysis method (Feature-based CNN) and CNN using images as input (Image CNN) to distinguish between chromosome and non-chromosome images. Beside, the prediction results from the above two methods are further combined to achieve higher classification accuracy (Hybrid CNN).

For an image feature analysis method, image recognition relies on the designed features. The discrimination power of the features would influence the classification results profoundly. In Feature-based CNN, we design seven features based on area, density, roughness and width of the images. The results indicate that these features are useful for distinguishing between chromosomes and non-chromosome objects.

The image features are designed manually and extracted from the images in Feature-based CNN. In addition, the image features are extracted via convolution and pooling process in Image CNN automatically. The information regarding image features in the two methods is obtained in different ways. As the feature information in Image CNN and Feature-based CNN is obtained independently, extra information should exist between the two methods. Based on this concept, we combine both prediction results from Image CNN and Feature-based CNN to improve classification performance. The results show that the combination of two prediction results has an effect on increasing the overall classification accuracy indeed. The combination of image-based and feature-based learning methods can improve classification performance.

Ninety-seven segmented objects, including 23 chromosomes and 74 non-chromosome objects, are mis-classified by the proposed method. Fig. 12 demonstrates some examples of major missing non-chromosome objects. On the other hand, small chromosomes with low density or chromosomes with highly curvature would be mis-classified as non-chromosome objects, as shown in Fig. 13. This phenomenon is the limitation of the proposed method.

**Fig. 12.**
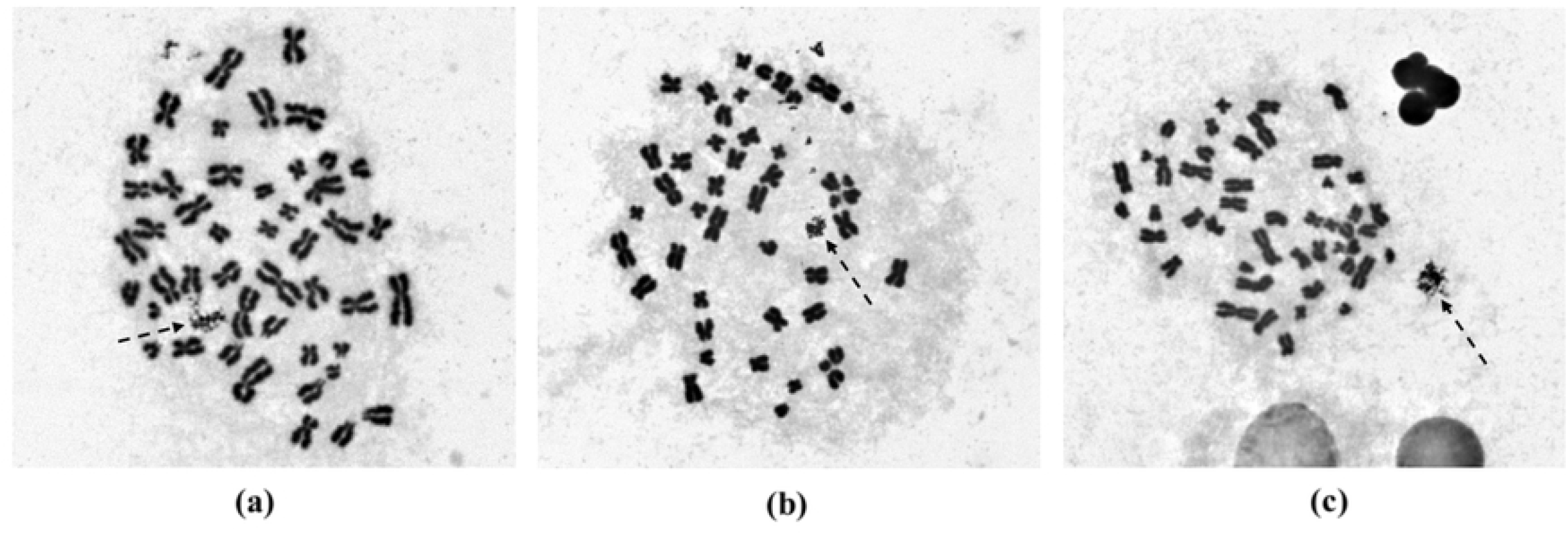
Non-chromosome objects indicated by arrows, which could not be detected and excluded by the proposed method.

**Fig. 13.**
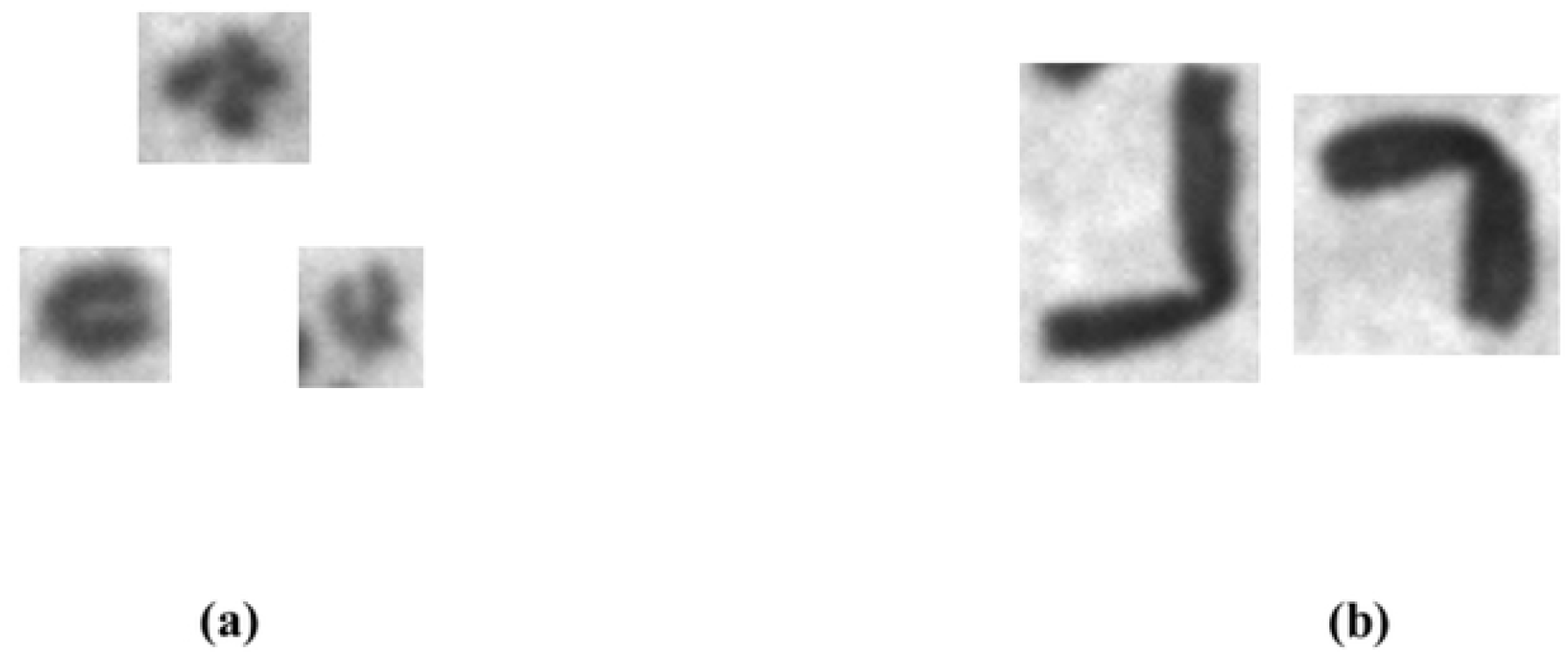
(a) small chromosomes with low density or (b) chromosomes with highly curvature would be mis-classified as non-chromosome objects by the proposed method.

Exclusion of non-chromosome objects is an essential preprocessing step in chromosome image analysis. In this study, we propose a hybrid deep learning method for detecting and excluding non-chromosome objects in chromosome images. The experimental results show that the proposed method can really be used in excluding non-chromosome objects and determining the chromosomes for further analysis.

## 5. Conclusions

We have proposed a novel method to exclude non-chromosome objects and preserve chromosomes for further analysis. In this paper, we use 1038 metaphase chromosome images to verify the performance of the proposed method. The experimental results show that the proposed method can filter out 99.61% (18,683/18,757) of the non-chromosome objects and keep 99.95% (43,375/43,398) of the chromosome objects. The proposed method has a high accuracy on excluding non-chromosome objects and could be used as a preprocessing procedure for chromosome image analysis.

## Acknowledgments

The authors wish to thank the Institute of Nuclear Energy Research for having providing chromosome images.

## Notes

### Competing Interest Statement

The authors have declared no competing interest.

